# What is the main driver of unsustainable natural resource use in the Comoro Islands?

**DOI:** 10.1101/2021.06.04.445177

**Authors:** Mohamed Thani Ibouroi, Said Ali Ousseni Dhurham, Aurélien Besnard, Nicolas Lescureux

## Abstract

The Comoros archipelago is a biodiversity hotspot by virtue of its high level of endemism. However, it suffers one of the highest rates of forest loss worldwide, mainly due to strong anthropogenic pressures. As Comorian populations depend on forest resources for subsistence, establishing relevant conservation strategies for their sustainable management requires the consideration of multiple stakeholders’ perspectives toward biodiversity and habitat conservation. To better understand the relationships between humans and nature; how comorian people use natural resource and the relevance of a protected area for long-term biodiversity conservation, we used Q-methodology to assess local people‘s perceptions regarding biodiversity and conservation actions. Three discourses are identified during analysis: “Pro-environment discourse”, “Keeping things as usual” and “Social and environmental concerns”. According to the results, employed respondents, were favorable to long-term forest and biodiversity conservation. In contrast, unemployed respondents were in favor of more immediate benefits while unemployed but educated respondents were in favor to both long-term forest conservation and immediate benefits from forests. This suggests that the lack of livelihoods for rural people is the main factor leading them to overharvest natural resources. These results suggest that biodiversity conservation of the Comoros archipelagos may benefit for plan aiming at (1) developing tourism and maintaining sustainable production of crops and livestock that could allow enhancing the livelihoods and well-being of all social groups, (2) developing projects such as local markets that could allow villagers to sell their agricultural production, (3) setting up awareness campaign for tree-planting and reforestation. Reforestation could allow re-establishing natural plants and make large trees available for long-term purposes.

## 1 INTRODUCTION

Biodiversity and natural resources provide many direct as well as indirect services to human society, including playing a crucial role in sustaining people‘s well-being (Giannini et al., 2012). As a consequence, human populations strongly depend on natural ecosystems (Zhu et al., 2016). This is especially true for the poorest populations of developing countries, who largely rely on wild plants for building materials and for natural medicines and food, and on wild animals for meat (Ryan et al., 2016). However, on a global scale, biodiversity and natural resources are being degraded at alarming levels, mainly induced by anthropogenic pressures (Brook et al., 2008).

Over the past two decades, scientists and numerous national and international organizations have argued for the urgent need to find alternative community-based approaches to protect and manage natural systems in developing countries (Jantz et al., 2015; King et al., 2021). Untill recently, natural resource and habitat management strategies tended to rely on biological and ecological data based on species ecology, population genetics or demographics, but have often neglected the human societies that critically depend on natural ecosystems (Fritz-Vietta, 2016, Gaebel et al., 2020; König et al., 2021). Although some conservation strategies have been developed in many countries on collaborative governance processes and participatory protected area management for instance, such strategies are non-existent in different parts of the world (Krueck et al., 2019; Ghosh-Harihar et al., 2019; Ayivor et al., 2020; Rittelmeyer 2020; O‘Brien et al. 2021; Jin et al., 2021; Arumugam et al., 2021). Communities living in geographic proximity to natural resources and forests typically have traditional knowledge about as well as emotional bonds with these areas. Ignoring the needs and practices of local communities in habitat conservation initiatives may result in conflicts between natural resource managers and these populations if the latter feel they face restrictions in the benefits they acquire from these areas (Fisher et al., 2020; Gaebel et al. 2020). This can eventually have a negative effect on both the long-term effectiveness of biodiversity conservation and on the livelihoods of the local population (Sournia, 1990; Fritz-Vietta, 2016; Debata et al., 2017; Gaebel et al., 2020; Jin et al., 2021). Reconciling the needs of the local population and natural resource use is now seen as fundamental in developing countries to implement management plans that ensure livelihoods and well-being in parallel with biodiversity conservation objectives (Helm Aveliina, 2006; Boron et al., 2016; Jin et al., 2021).

The Comoros (an archipelago consisting of the islands of Anjouan, Grande Comore, Mohéli and Mayotte) is a biodiversity hotspot by virtue of its high level of endemism (Myers et al., 2000). However, on the islands of the Union of Comoros (Grande Comore, Anjouan and Mohéli), natural habitats are experiencing one of the highest rates of habitat loss in the world (9.3% each year, FAO, 2010). The Union of Comoros is also one of the poorest nations in the world (Bourgoin et al., 2017). According to Fisher and Christopher (2007), about 72% of Comorians depend directly on forest resources for subsistence (Fisher and Christopher, 2007; Bourgoin et al., 2017). Some 60% of Comorians live below the poverty line and 49% are undernourished. Additionally, the Union of Comoros has a fast-growing population, leading to an acute need of land for agriculture and wood for building (Elvidge et al., 2009). Many researchers have pointed to intensive land use as the direct cause of the very high rate of natural habitat loss observed in the archipelago (Ibouroi et al., 2018a, b). Yet this pressure on natural forests and biodiversity is altering the ecosystem services they provide for the Comorian people. Effective conservation strategies are crucially needed to ensure the long-term preservation of biodiversity and natural habitats in the Comoros.

On the three islands of the Union of Comoros, some measures have been undertaken by local, national, and international organizations in the aim of ensuring the long-term conservation of biodiversity (Granek and Brown, 2005; Poonian et al., 2008; Ibouroi et al., 2018b; Ibouroi et al., 2019). For instance, in 1992, Mickleburgh et al. proposed a long-term monitoring of the Livingstone‘s flying fox population and the establishment of a captive-breeding program for the species (Mickleburgh et al., 1992). The creation of the Mohéli Marine Park was successful in 2001 (Granek and Brown, 2005). Some of these projects were funded by the United Nations Development Program (UNDP 1998). In 2016, the national network of marine and terrestrial protected areas was created in the three islands of the Union of Comoros (see Ibouroi et al., 2019). However, most of these conservation strategies have been restricted to protecting Livingstone‘s flying fox roosts (Ibouroi et al., 2018b), as this is one of the most endangered species on the islands. Strategies to conserve the islands’ biodiversity and habitats need to consider various contentious aspects that currently involve complex decision-making dilemmas (e.g. forest management, hunting management, representation of local communites, etc.). Solutions have not yet been clearly defined. For instance, numerous gaps still remain in understanding stakeholders’ perspectives regarding natural resource management and biodiversity conservation. Local people‘s subjectivity and viewpoints are important to identify in order to inform conservation strategies and future management practices, to avoid making mistaken decisions in planning these measures and to increase their chance of being effective (Niedziałkowski et al., 2018).

In this study, we conducted a Q-methodology approach to assess the relationships between stakeholders and their use of natural resources as well as their impact on habitats in the Comoros. Specifically, we assessed (1) how stakeholders perceive benefits from natural resources, (2) the level of awareness of the impact of their practices on biodiversity, and (3) their knowledge about, perceptions of and attitudes toward biodiversity and conservation actions. As social factors such as the level of formal education, employment and geographic location can affect knowledge and determine attitudes, we assessed what factors were related to positive or negative perception of forests and biodiversity conservation. This information may help (1) to understand the local community‘s representation of biodiversity, and (2) to explore future scenarios, with the objective of proposing relevant long-term conservation actions and habitat management strategies.

## 2 MATERIALS AND METHODS

### 2.1. Study area

The Comoros archipelago is located in the Indian Ocean, midway between Madagascar and the eastern coast of Africa. This archipelago comprises four islands: Grande Comore, Mohéli, Anjouan (the Union of the Comoros), and Mayotte (an overseas department of France). Without Mayotte, the Comoro Islands cover 1,862 km^2^ and represent the third smallest African nation in terms of surface area (Michon, 2016). The islands are separated from each other by a distance of about 40–80 km. Since their emergence about 7 million years ago, these islands have never been connected to a continental mainland or to each other (Louette et al., 2004). Our study focused specifically on the three islands of the Union of the Comoros.

In the Union of Comoros, habitat fragmentation and loss differ between islands due to differences in habitats, ecology and human demographics among islands (Sewall et al., 2007). For example, Anjouan Island experiences the highest human population density (772.13 inhabitant/km^2^ against 180.55 inhabitant/km^2^ and 357.78 inhabitant /km^2^ for respectively Mohéli and Grande Comoro Islands) within the archipelago, which has direct consequences on natural habitat disturbance. On this island, between 1972 and 1987, more than 85% of natural habitat was converted into farmland, urban areas and secondary forests (Goodman et al., 2010). In the Grande Comoro Island, the rate of habitat loss is also high but in certain regions for instance in the Karthala forest, habitat fragmentation is moderate. In contrast, both habitat loss and fragmentation are relatively limited on Mohéli, probably because of the presence of a protected area (the Mohéli Marine Park) but also due to the low human population density on this island (180.55 inhabitant/km^2)^. The Mohéli Marine Park was established in 2000 with the goal of protecting 404 km^2^ of marine habitats home to many endemic and threatened taxa, such as the dugong (*Dugong dugon*) and the green sea turtle (*Chelonia mydas*). The presence of this marine protected area represents an important source of income for local communities – several members of the community have been hired by the park as regular staff (Granek and Brown, 2005). Many tourists also come to see the endemic marine taxa and then take the opportunity to discover endangered terrestrial species such as the Livingstone‘s flying fox (*Pteropus livingstonii*) and the mongoose lemur (*Eulemur mongoz*). This tourist activity generates direct incomes for some local people (for example, who work as guides or in hotels, etc.). Our study involved different localities on the three islands of Comoros (Anjouan, Mohéli and Grande Comore, Fig 1).

**Figure 1:**
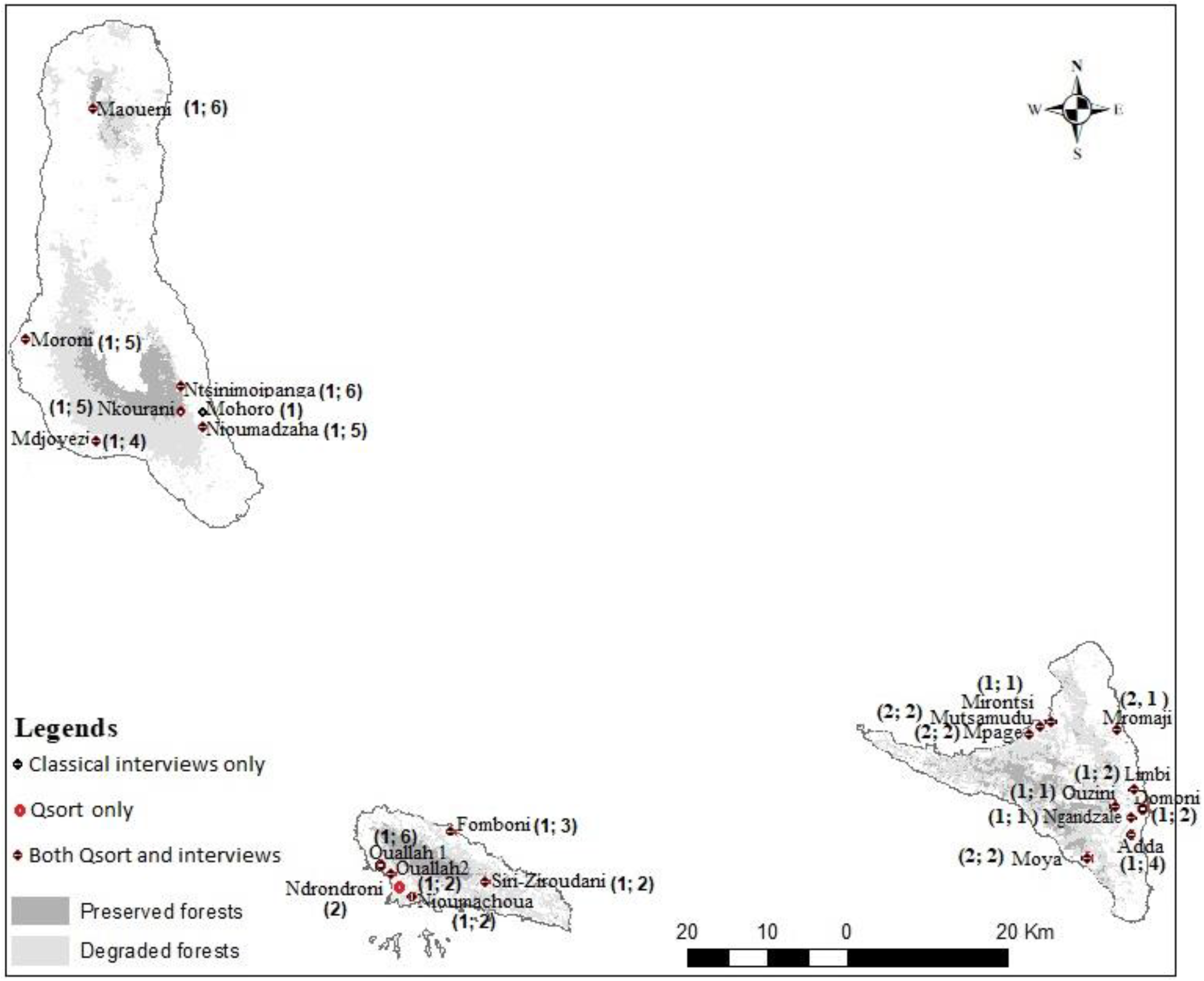
Sampling localities in the Union of Comoros: Grande Comore (top left), Anjouan (bottom right), Mohéli (bottom left), indicating main villages and preserved (dark grey) and degraded (light grey) forests; Numbers of interviewed people are presented between brackets for each locality; when both classical interviews and Qsort methods are realized in the same locality, the first number on the parentheses represents the number of interviewed people for classical interviews and the second represents the number for Qsort.

To understand how stakeholders perceive benefits from natural resources, their knowledge, perceptions and attitudes toward biodiversity and conservation actions, some of the interview questions and Q statements centered on two native flying fox species: Livingstone‘s flying fox (*Pteropus livingstonii*) and the Seychelles fruit bat (*P.seychellensis comorensis*), which differ in their feeding and roosting behavior as well as in their dispersal patterns (Norberg et al., 2000; Ibouroi et al., 2018a). *Pteropus livingstonii* is confined to the remaining mountain forests on Anjouan and Mohéli and feeds on endemic forest plants, while *P. s. comorensis* is widely distributed over the four islands of Comoros, feeding in both forests and cultivated areas (Ibouroi et al., 2018b; Trewhella et al., 2001). Both species are important ecosystem service providers, as they are pollinators and seed dispersers (Ibouroi et al., 2018b). Their differences in habitat use and feeding ecology ensure different ecosystem services. The two species have a potentially crucial impact on both forest regeneration and the cultivation of crops, thus are critical for maintaining overall ecosystem dynamics (Ibouroi et al., 2018b). Because of this contrasted pattern of dispersal, feeding and roosting behavior, conservation strategies and conflicts between humans and bats are also different between the two species. For instance *Pteropus livingstonii* populations are the subject of conservation actions, some of which involve local communities. These conservation actions focus on this species not only because of its low population size but also the rapid forest loss in the Comoros (Ibouroi et al. 2018b). Regarding *P.seychellensis comorensis*, as the species roosts and feeds in overexploited forests, its population is commonly involved in conflicts, as individuals feed in farmed areas and can damage cultivated plants. Such conflicts are believed to be the primary driver of legal and illegal persecution of this species, as is the case in many countries (Oleksy et al., 2018). For these reasons, these species are an ideal model to investigate local Comorian perceptions, allowing their discourses to be mapped regarding the flying fox, biodiversity and social development, followed by an analysis of the consequences for the long-term conservation of natural habitats.”

### 2.2. Research Design

Q-methodology is a standard method used to reveal people‘s subjectivity and explore viewpoints on defined issues that are often contested (Stephenson, 1935). It specifically aims at identifying underlying patterns among stakeholders and comparing the key viewpoints, which leads to the identification of shared broad common points as well as divergences between them (Watts and Stenner 2005, 2012; Bavin et al.,2020; Arumugam et al., 2021). The approach combines the qualitative study of attitudes with the statistical rigors of quantitative research techniques (Watts and Stenner, 2005, 2012; Bavin et al.,2020; Arumugam et al., 2021). It is increasingly applied in different types of environmental research, including environmental management and policy and social science of conservation (Debata et al., 2017; Kamal and Grodzinska-Jurczak, 2014; Niedziałkowski et al., 2018; Walder and Kantelhardt, 2018, Rittelmeyer 2020; Fisher et al., 2020; Arumugam et al., 2021).

Q-methodology involves five main steps: (1) collecting a broad sample of statements (**concourse and Q-set design**); (2) Selecting a representative sample of statements (reflecting the diversity of the wider concourse) to consider as Q-set (‘**Formulating the Q-Set**’); (3) Selection of participants (‘**Identifying the P-Set**’); (4) Conducting the Q-sorts and Interviews (‘**Q sorting and post-sorting interview**’); (5) Analyzing the data using factorial analysis (‘**Analyzing the data and development of factor perceptions**’) (Eden et al., 2005; Kamal and Grodzinska-Jurczak, 2014). The standardized steps of Q methodology are summarized in the Figure 2.

**Figure 2.**
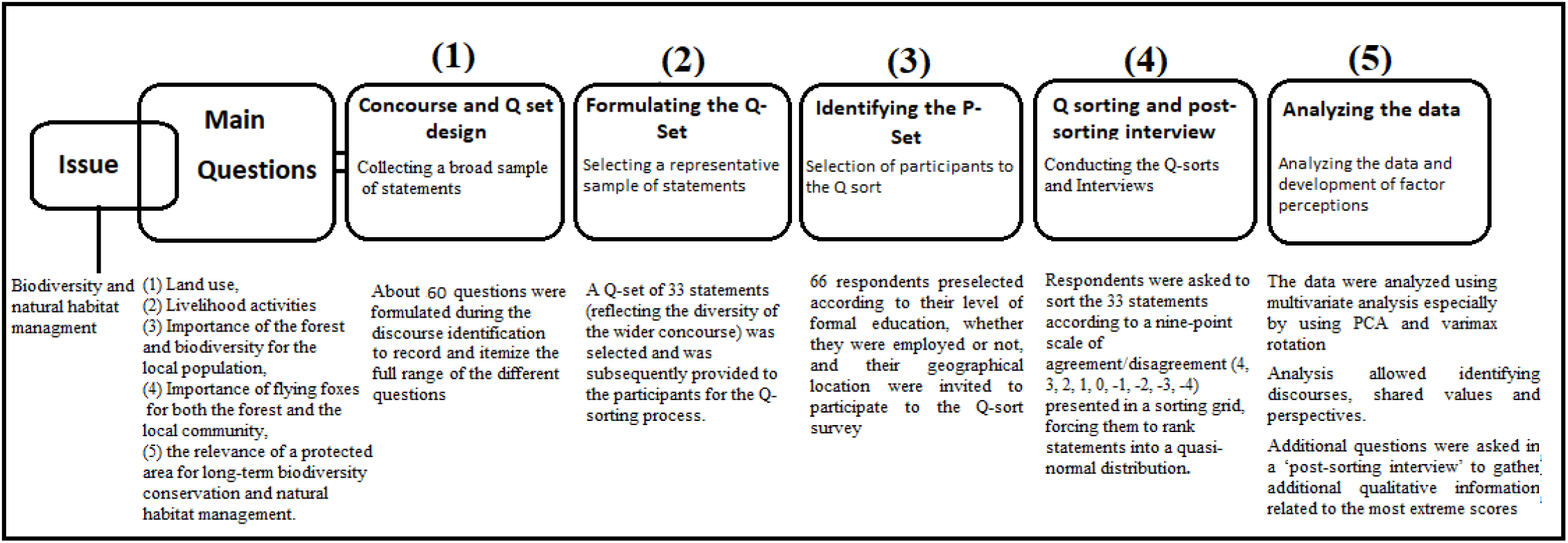
Q methodology steps conducted for the current study.

#### 2.2.1 Concourse and Q set design

Q-methodology was conducted in three phases: between August and October 2016, January and April 2018, and between December 2018 and March 2019. In each field session, the three islands were visited for collecting data. As a first step, we established a concourse, defined as the full opinion spectrum in relation to the topic of habitat and natural resource uses, biodiversity and habitat conservation. For this concourse establishment, we used semi-structured interviews with local population during our first field session (August and October 2016) to gather information regarding the forest and natural resource uses, land uses and biodiversity conservation. These semi-structured interviews were based on pre-defined interview questions (Table 1) and all the people interviewed were not preselected but directly asked to participate to the interview when encountered in villages in the course of their daily activities or during our prospection in forests. Each discussion and the recording of the collected information took about one hour. All responses were recorded with a dictaphone. For each person interviewed, their gender, age, place of residence, socio-professional activity, and level of formal education were recorded. In total, 40 people were asked to participate in the interviews, of which 13 (1 man and 12 women) declined and 27 agreed. Of the 27 people interviewed, one respondent was under the age of 18 and was excluded from the analysis. The other 26 respondents were 23 men and 3 women aged between 22 to 65 (average age of 41); 14 lived on the island of Anjouan, 7 on Mohéli, and 5 on Grande Comore. From the final interview transcripts, a total of 60 statements were extracted to form the concourse.

**Table 1.**
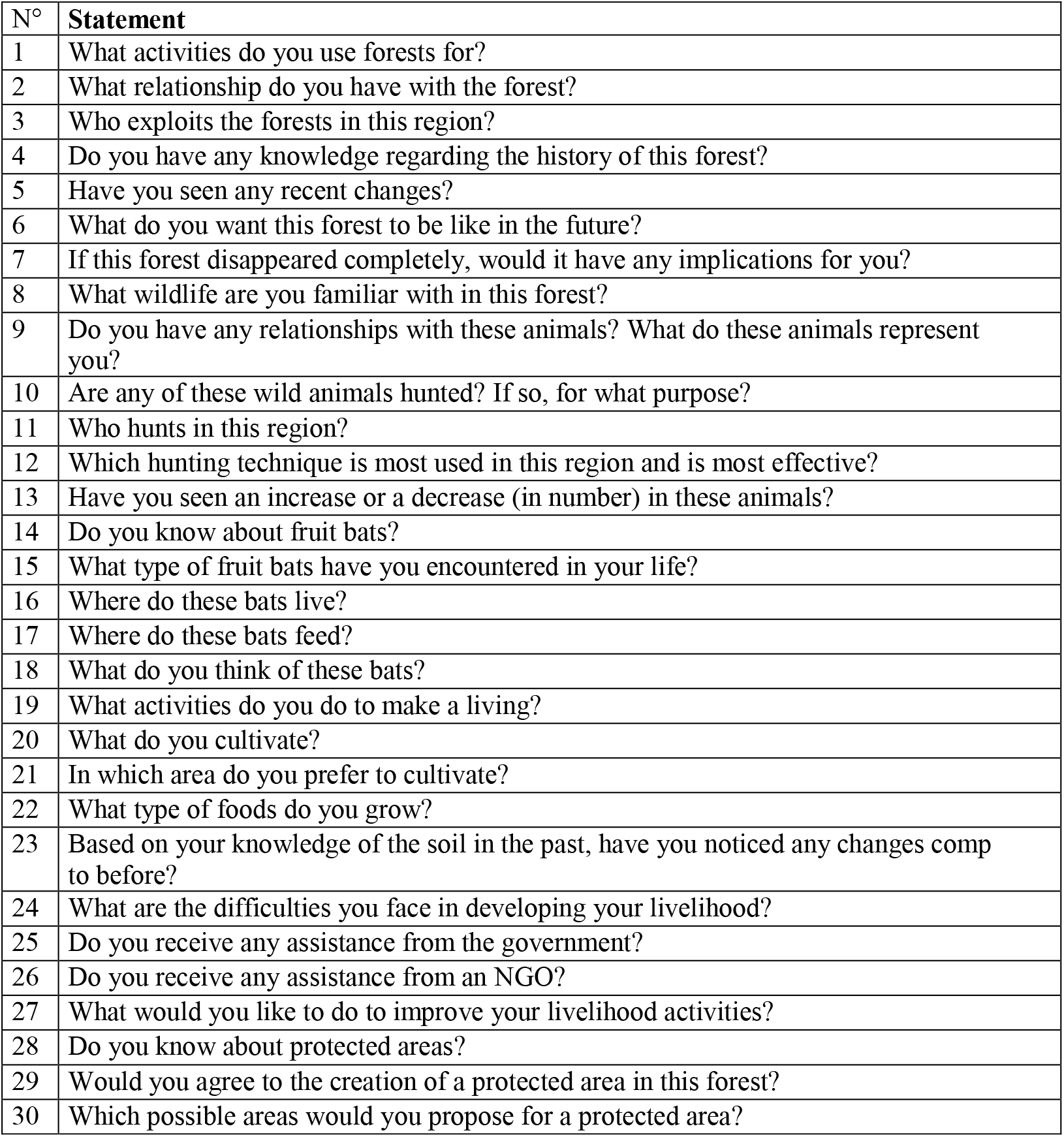
Guidelines for the initial semi-structured interviews with rural Comorians

#### 2.2.2 Formulating the Q-Set

Within the 60 statements selected as concourse (see above), a final set of 33 statements (Figure 2) were selected as ‘Q-set’ by using a structured filtering process in order to reduce the whole concourse into a manageable set of statements. Statements expressing the same value or viewpoints were summarized into one overarching statement. These 33 statements (Q-set or Q-sort, Fig 3) representing the diversity of the wider concourse cover five main topics:(1) land use, (2) the livelihood activity of the local population, (3) the importance of the forest and biodiversity for the local population, (4) the importance of flying foxes for both the forest and the local community, and (5) the relevance of a protected area for long-term biodiversity conservation and natural habitat management.

**Figure 3:**
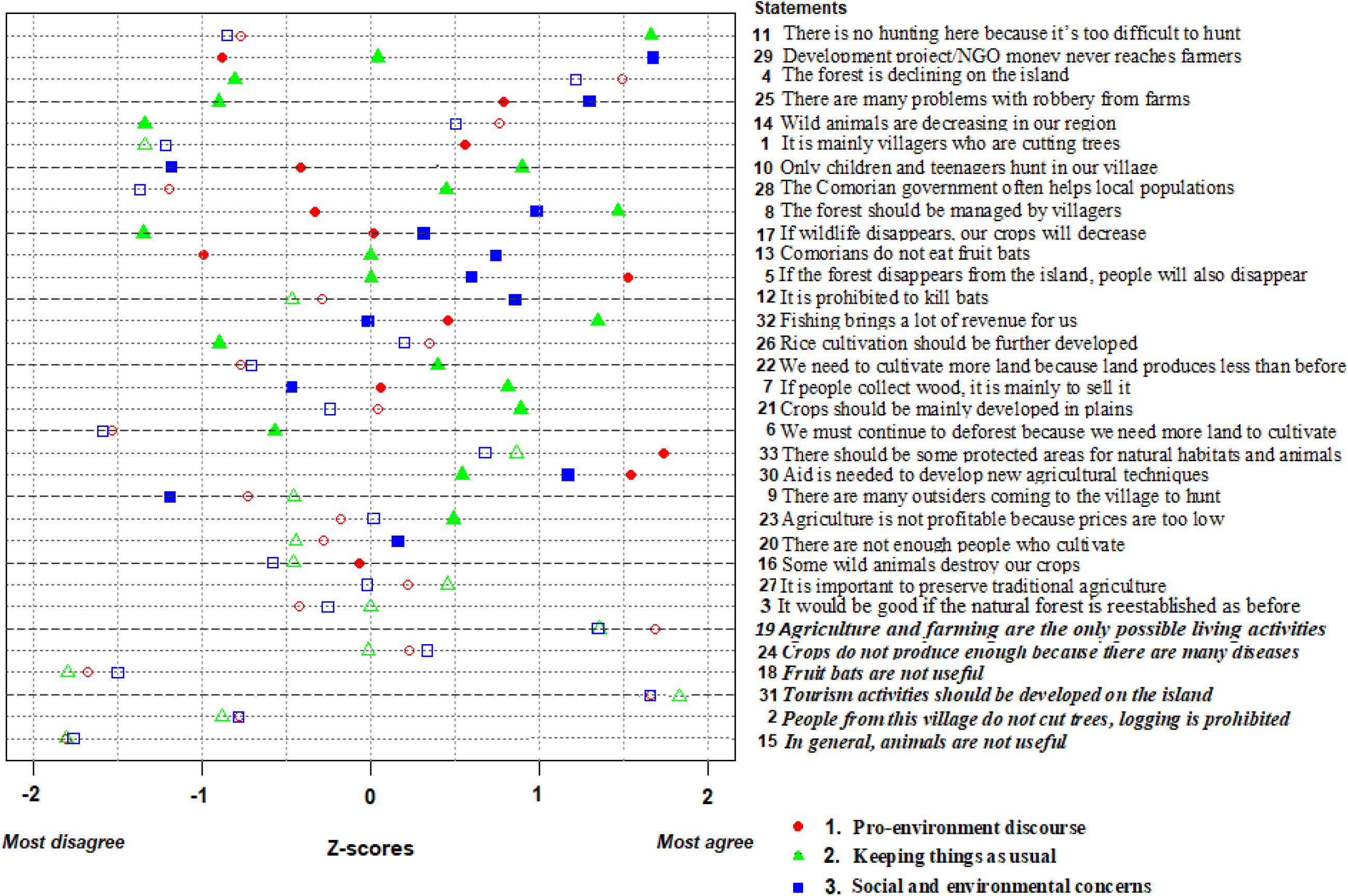
Statements selected for Q sorting, ordered from most distinctive (top) to consensus (down, in bold and italic), based on Z-score differences; A statement is considered distinctive when comparing all pair of factors and at least one factor is significantly different to the others for this statement at p-value < .01 (e.g. statement 11); if all the comparisons between each pair of factors are significantly different at p-value < .01, the statement is considered as “distinctive all” (e.g. statement 29); a statement is considered as consensus when none of the comparisons are significantly different at p-value < .01(e.g. statement 15); if a statement is distinctive for a factor (at p < 0.01), the symbol is filled and if a statements is not distinctive for a given factor, the symbol is empty.

#### 2.2.3 Identifying the P-Set

Typically, Q methodology involves a relatively small number of respondents, varying from 26 to 46 (Zabala et al., 2018) although some few studies used large number of respondents beyond 100 individuals (Milcu et al., 2014; Carmenta et al. 2017; Zabala et al., 2018). Although respondents involved in Q-study have to be diverse, the sample does not have to be representative of the population as the aim is to get the most diverse range of opinions, regardless of whether they are minority ones (Zabala et al., 2014). In order to represent a range of opinions from local people, 66 respondents (P-set, 51 men [77%] and 15 women [23%]) who had not participated in the previous semi-structured interviews were invited to complete the sorting and post-sorting interview. In contrast to the concourse stage, Q-sort respondents were firstly preselected according to their level of formal education, whether they were employed or not, and their geographical location. This selection was firstly based on our knowledge in the Comoros institutions and forest workers, local networks and collaboration but also based on a snowball sampling approach (i.e. the identification of stakeholders by other participants). In the field, we get other information regarding villagers working in conservation and environmental institutions/NGO but also villagers with high/low level of education for each locality. These villagers are selected as respondents for the Q-Method process.

#### 2.3.4 Q sorting and post-sorting interview’

In the Q-sorting and post-sorting process, a researcher presents the statements (Q-Set) so participants (P-Set) can rank them according to the predefined Q-sort structure in order to express their level of agreement or disagreement with. Interviews were conducted on a face-to-face. As for the semi-structured interviews, discussion and the recording of the collected information took about one hour and interviews were conducted in local language. For each respondent, the gender, age, place of residence, socio-professional activity, and level of formal education were also recorded. The researcher had to explain to all participants that the aim of the Q-sorting process was to obtain their opinions rather than to test for their knowledge. Participants represented by men and women but also by people from urban vs. non-urban regions (see table 2) were given the Q-set and were instructed to read the statements carefully. They were asked to sort the 33 statements according to a nine-point scale of agreement/disagreement (4, 3, 2, 1, 0, −1, −2, −3, −4) presented in a sorting grid, forcing them to rank statements into a quasi-normal distribution (see Fig s1). Each participant was then asked to explain their most extreme scores (−4 and +4), and these comments were later used to interpret the results.

**Table 2:** Factor matrix based on Q methodology (* indicates that the corresponding respondent loaded to a narrative), Emp= Employed outside NGOs; EmpNGO= Employed in an NGO; G.Comoro= Grande Comore; Educ= educated; Non-educ= non-educated; the 5 respondents that didn’t fit any of the 3 narratives are shown in bold; M=Male; F= Female

| Island | Gender | Age | Urban/non-urban | Education | Social group | Narrative A | Narrative B | Narrative C |
| --- | --- | --- | --- | --- | --- | --- | --- | --- |
| G.Comore | F | 26 | Non-urban | Educ | Unp | 0.19 | * 0.40 | 0.13 |
| Mohéli | M | 19 | Non-urban | Non-educ | Unp | 0.22 | -0.14 | * 0.54 |
| Mohéli | M | 20 | Non-urban | Non-educ | Unp | * 0.36 | 0.33 | 0.03 |
| Mohéli | M | 45 | Non-urban | Educ | Emp | * 0.63 | 0.27 | 0.21 |
| Mohéli | M | 45 | Non-urban | Non-educ | Unp | -0.08 | 0.31 | * 0.37 |
| Mohéli | M | 65 | Non-urban | Educ | Emp | 0.54 | 0.14 | * 0.64 |
| Mohéli | M | 26 | Non-urban | Educ | Emp | * 0.58 | 0.04 | 0.14 |
| Mohéli | M | 36 | Non-urban | Non-educ | Unp | 0.04 | 0.23 | * 0.46 |
| Mohéli | M | 41 | Non-urban | Educ | Emp | * 0.60 | 0.14 | 0.47 |
| **Mohéli** | **F** | **56** | **Non-urban** | **Educ** | **Emp** | **0.41** | **0.42** | **0.38** |
| Mohéli | M | 33 | Non-urban | Educ | Unp | 0.42 | -0.06 | * 0.51 |
| Mohéli | M | 26 | Non-urban | Educ | Emp | * 0.62 | 0.13 | 0.34 |
| Mohéli | M | 37 | Non-urban | Educ | Emp | * 0.76 | 0.16 | 0.34 |
| Mohéli | M | 29 | Non-urban | Educ | Emp | * 0.54 | -0.14 | 0.24 |
| Mohéli | M | 46 | Non-urban | Educ | Emp | * 0.57 | -0.15 | 0.29 |
| G.Comore | M | 58 | Non-urban | Educ | Emp | * 0.72 | -0.01 | 0.35 |
| Anjouan | M | 53 | Non-urban | Educ | Emp | * 0.56 | 0.10 | 0.39 |
| Anjouan | M | 42 | Non-urban | Educ | Emp | * 0.58 | 0.11 | 0.53 |
| Anjouan | M | 52 | Urban | Educ | Emp | 0.36 | 0.25 | * 0.60 |
| Anjouan | M | 37 | Urban | Educ | Emp | * 0.58 | -0.05 | 0.46 |
| Anjouan | M | 31 | Non-urban | Educ | Emp | * 0.69 | 0.28 | 0.39 |
| Anjouan | M | 27 | Non-urban | Educ | Emp | * 0.71 | 0.08 | 0.21 |
| Anjouan | M | 45 | Non-urban | Educ | Emp | * 0.69 | 0.14 | 0.40 |
| Anjouan | M | 38 | Non-urban | Non-educ | Unp | 0.32 | 0.33 | * 0.51 |
| **Anjouan** | **M** | **25** | **Non-urban** | **Educ** | **Unp** | **0.39** | **0.38** | **0.39** |
| G.Comore | M | 47 | Non-urban | Educ | Emp | * 0.80 | 0.21 | 0.25 |
| G.Comore | M | 23 | Non-urban | Educ | Unp | 0.05 | 0.20 | * 0.39 |
| G.Comore | M | 33 | Non-urban | Non-educ | Unp | * 0.66 | -0.07 | 0.25 |
| G.Comore | M | 35 | Non-urban | Non-educ | Unp | 0.18 | 0.08 | * 0.75 |
| G.Comore | M | 36 | Non-urban | Non-educ | Unp | 0.20 | 0.12 | * 0.35 |
| Anjouan | F | 25 | Urban | Educ | EmpNGO | * 0.71 | 0.10 | 0.15 |
| Anjouan | F | 24 | Urban | Non-educ | Unp | 0.14 | 0.16 | * -0.42 |
| Anjouan | M | 27 | Non-urban | Educ | EmpNGO | * 0.54 | 0.25 | 0.16 |
| G.Comore | M | 40 | Non-urban | Non-educ | Unp | 0.12 | * 0.96 | 0.12 |
| Anjouan | F | 26 | Urban | Educ | EmpNGO | * 0.81 | 0.24 | 0.18 |
| Anjouan | F | 22 | Urban | Non-educ | Unp | 0.50 | * 0.60 | -0.01 |
| **Anjouan** | **M** | **38** | **Urban** | **Educ** | **EmpNGO** | **0.42** | **0.20** | **0.43** |
| Mohéli | F | 28 | Urban | Non-educ | Unp | * 0.63 | 0.16 | 0.28 |
| Mohéli | F | 35 | Urban | Non-educ | Unp | 0.23 | 0.17 | * 0.52 |
| Mohéli | M | 42 | Urban | Non-educ | Unp | * 0.52 | -0.04 | 0.12 |
| G.Comore | M | 38 | Non-urban | Non-educ | Unp | 0.08 | * 0.98 | 0.09 |
| G.Comore | F | 42 | Non-urban | Non-educ | Unp | 0.12 | * 0.96 | 0.12 |
| G.Comore | F | 26 | Non-urban | Educ | EmpNGO | 0.25 | 0.08 | * 0.70 |
| G.Comore | M | 35 | Urban | Educ | EmpNGO | * 0.65 | -0.06 | 0.39 |
| G.Comore | M | 44 | Non-urban | Educ | EmpNGO | * 0.52 | 0.33 | -0.31 |
| G.Comore | M | 29 | Non-urban | Non-educ | Unp | 0.15 | * 0.55 | 0.05 |
| **G.Comore** | **M** | **35** | **Non-urban** | **Non-educ** | **Unp** | **0.40** | **-0.36** | **0.48** |
| G.Comore | M | 26 | Urban | Educ | EmpNGO | * 0.74 | 0.14 | 0.24 |
| G.Comore | F | 37 | Non-urban | Non-educ | Unp | 0.08 | * 0.98 | 0.12 |
| G.Comore | M | 36 | Urban | Educ | EmpNGO | * 0.44 | 0.28 | 0.13 |
| G.Comore | M | 29 | Urban | Educ | EmpNGO | * 0.66 | 0.12 | -0.24 |
| G.Comore | M | 61 | Urban | Educ | EmpNGO | 0.52 | 0.16 | * 0.57 |
| G.Comore | M | 42 | Urban | Educ | EmpNGO | * 0.77 | 0.19 | 0.14 |
| Anjouan | M | 38 | Urban | Educ | EmpNGO | * 0.78 | 0.18 | 0.16 |
| G.Comore | F | 40 | Non-urban | Non-educ | Unp | 0.03 | * 0.96 | 0.17 |
| Anjouan | M | 42 | Urban | Educ | EmpNGO | * 0.73 | 0.28 | 0.12 |
| G.Comore | M | 32 | Urban | Educ | Emp | 0.40 | -0.03 | * 0.48 |
| G.Comore | M | 35 | Urban | Educ | EmpNGO | * 0.57 | 0.13 | 0.07 |
| Anjouan | M | 37 | Urban | Educ | EmpNGO | * 0.85 | 0.14 | 0.24 |
| **G.Comore** | **M** | **42** | **Urban** | **Educ** | **EmpNGO** | **0.41** | **0.25** | **0.42** |
| G.Comore | F | 39 | Non-urban | Non-educ | Unp | 0.08 | * 0.98 | 0.09 |
| G.Comore | M | 38 | Non-urban | Educ | EmpNGO | * 0.55 | 0.15 | 0.49 |
| G.Comore | M | 30 | Urban | Educ | Unp | 0.35 | 0.19 | * 0.45 |
| G.Comore | F | 42 | Urban | Educ | EmpNGO | * 0.83 | 0.13 | 0.16 |
| G.Comore7 | F | 32 | Non-urban | Educ | EmpNGO | * 0.77 | 0.17 | 0.15 |
| G.Comore | M | 37 | Non-urban | Non-educ | Unp | 0.10 | * 0.98 | 0.10 |

During this Q-sort process, some of the difficulties encountered were: (1) The fact that the method is time-consuming in the preparation, data collection, and analysis phases. For instance: (a) because of the high rate of poverty in the Comoros, a large number of our potential participants, especially those working in forests, declined to participate unless they were paid. (b) Our semi-structured and Q-sort sampling involved only a small number of woman‘s as they tended to decline to be interviewed probably for reasons related to the local culture. The few interviewed women were mainly employed in NGOs and students. No woman met in villages agreed to be interviewed. This is probably because the pre-selection of participants from the different villages were carried out few days before the interviews and no discussion was made with their husbands or legal parents. (2)Because respondents were often selected few days before the Q-sorting process, some of them did not have basic knowledge of the questions and often answered haphazardly. This that can impact our results as the goal of the research is to use a set of relevant people and a sample of opinion statements to draw conclusions.‖

#### 2.3.5 Analyzing the data and development of factor perceptions

The data were analyzed using the ‘qmethod’ package for R (R Development Core Team, 2016; Zabala, 2014) which groups responses according to their similarity, using PCA and varimax rotation (a common approach in Q methodology). Different factors were rotated and compared during the multivariate analyses. We choose three factors based on a combination of total explained variance, minimum correlations between factors and reduced number of confounders (participants loading on more than one factor). These three factors were retained as different discourses because they had the minimum of two or more significantly loading participants (at p < 0.01 level, threshold value = 2.58 *1/ √ (number of statements = 33) = ± 0.44).”

Additionally, we analyzed the dataset with an inter-class Principal Component Analysis (PCA) implemented in the ade4 R package (Thioulouse and Dray, 2007) in order to easily identify contrasted statements between the different social groups. This method which doesn‘t require parametric data (it is not based on any probabilistic model, but only on geometric considerations) rotates the selected PCA axes to maximize correlation between predefined groups. In a first analysis, we tested the discrimination between (1) the group of employed people working in NGOs (EmpNGO), (2) the group of employed people not working in NGOs (Emp), and (3) the group of unemployed people with a low level of formal education (unp). In addition, we tested the discrimination between (4) people from the three islands of the archipelago, (5) people from urban vs. non-urban regions, (6) age classes (ages were classifed as young [18 to 35 years] and old [36 to 75 years]) and (7) the gender (men and women groups). We tested whether these predefined groups significantly differed from each other in terms of Q-sort scoring using a permutation test based on 1000 permutations. The tests were considered significant when the p-value was < 0.05.”

## 3. RESULTS

### 3.1. Semi-structured interview responses

All 26 people interviewed stated that they receive benefits from forests and use natural resources for everyday life. All respondents stipulated that they go to the forest to work (for agriculture and cultivation and to collect wood). In answer to the question “If this forest disappears completely, would that result in changes for you? What influence does the forest have on your well-being?”, most respondents highlighted that the forest is essential for fertile soil, and thus necessary for agriculture, and is also important in maintaining the water source.

A large majority of the respondents stated that they know what biodiversity is and its importance for their subsistence and most of these have a positive perception of wild animals and reported that these are useful for their well-being. Only a few minority of our respondents stated that some wild animals are harmful. Comorian attitudes toward bats were mostly positive and only a minority reported that they did not know the usefulness of fruit bats. Of those with a positive attitude and perception of fruit bats, most reported their importance (1) as seed dispersers for forest regeneration, (2) as seed dispersers for important cash crops such as cashews and mangos, (3) as pollinators, or (4) as a source of income from tourism (the case of *P.livingstonii*). Some respondents mentioned that fruit bats, especially those living in villages (*P.s.comorensis*), generate some damage in cultivated areas.

All the interviewees had some knowledge about the primary forest and its usefulness for the local population. In answer to the question “Have you noticed any recent changes?” regarding their perception of landscape changes within the forest, a large majority reported that the forest is overharvested and is decreasing in surface area. They highlighted that the decline of the forest is having an impact on their livelihood. When asked “Who exploits the forests in this region?”, they gave contrasting responses. Some respondent reported that villagers are responsible for forest loss due to the practice of intensive wood collection and only a minority of respondent claimed that their forests are harvested by foreigners from other cities on the island.

Despite these diverging views, all respondents reported the negative impact of forest misuse on their livelihoods and well-being, and stated that if forests disappear completely, human life will not be possible in their region.

Most respondents reported that rural populations are neglected and lack assistance from the government and/or NGOs, stating that this is the main cause leading to forest overharvesting. A large majority reported that they never benefit from any government assistance or help from NGOs and said that the lack of agricultural equipment and technical assistance are the main factors inciting rural people to harvest the forest. They mentioned that the lack of assistance from the government and NGOs forces the rural population to be highly dependent on forests as they do not have any alternative livelihood. On this point, the local population agreed that forests must be protected or even regenerated and a majority agreed with the creation of protected areas in their region, and only a minority agreed under certain conditions, notably governmental support of their livelihoods and for agricultural equipment and technology.

### 3.2. Q sorting and post-sorting results

Among the 66 respondents interviewed during the Q-sort possess, five individuals (four men and one woman, see table 2 for age and demographic repartition) did not agree with any discourse as they had low sorts loadings on all factors thus were considered non-significant at p < 0.01.

Of the 33 Q statements (see Fig 3), six (18%) were consensual for all respondents (either positive or negative) and thus did not contribute to discriminations in discourse. Altogether, these discourses explained 55% of total variance. These three discourses were labeled according to the different statements significantly loaded to the considered factor (narrative A: “Pro-environment discourse”, narrative B: “Keeping things as usual”, and narrative C: “Social and environmental concerns”). The results found a low correlation between narratives A and B (r=0.30) and between narratives B and C (r=0.30), indicating that they are distinct (Table s1). The correlation between narratives A and C was higher (0.68), indicating some similarities between them (Table s1).

#### 3.2.1 Consensus statements

There was consensus on the need to develop tourism activities on the islands [statement 31], for example, all respondents agreed that “Tourism is important for Comoros development”. One respondent ranked this on the extreme end of the scale of agreement (+4) and commented: “We need to develop tourism; this is part of our development program in the Mohéli Marine Park” (see Fig 3). All respondents disagreed with the idea that animals and fruit bats are not useful [statements 15 and 18]. One respondent who strongly disagreed (−4) commented that “Animals are very useful; they represent food for local people and are very important for both forests and plantations”. Many responded in line with the comment, “Fruit bats are useful for ecotourism, for improving crops and the development of forests”. All respondents were rather neutral concerning the statement that agriculture and farming are the only possible livelihood activities on the island [statement 19], and there was general consensus, with slight disagreement, on the fact that people from the village do not cut trees as logging is prohibited [statement 2].

#### 3.2.2 Narrative A: Pro-environment discourse

Narrative A (factor 1) explained 27% of the total variance (Table s1). For this narrative, 35 of the 66 respondents loaded significantly (12 respondents from Anjouan, 10 respondents from Moheli and 13 respondents from Grande Comoro). These respondents were mainly employed, either in NGOs or in another sector (EmpNGOs =16 and Emp =15, see Table 2). They agreed with the statement that people will disappear from the islands if the forest disappears [statement 5; Factor 1 score: +3]. As one respondent commented, “The forest is our life: when it disappears from the island, we cannot survive.” Another participant who strongly agreed (+4) stated, “The forest is very valuable to our lives, if it disappears it will be catastrophic and will be the end of our lives.” Respondents in line with this narrative agreed with the fact that it would be good to reestablish the forest as before [statement 3; Factor 1 score: +4] and to have protected areas for habitats and animals [statement 33; Factor 1 score: +4]. For example, one respondent who strongly agreed commented, “It would be good to reestablish the forest as it was before. There used to be a diversity of foods, many rivers and it was wetter.” Another strongly agreeing respondent (+4) highlighted that “Dense natural forests are important; before, the forest brought more benefits than now.” They disagreed with the statement “There is a need to cultivate more land [statement 22]”. Instead, they agreed that “It is important to develop new agricultural techniques” [statement 30; Factor 1 score: +3]. As one respondent commented, “We need new methods and techniques to improve lands for cultivation that will allow us to increase production.” Other comments included: “We need materials and methods for agriculture that are more ecological.” “Technical and material aid is important as this will allow us to improve agricultural production.” Those associated with this narrative disagreed with the fact that Comorians do not eat fruit bats (Table 3). One respondent affirmed, “Some Comorians eat fruit bats, I can confirm this as I have been present in many cases.”

**Table 3:**
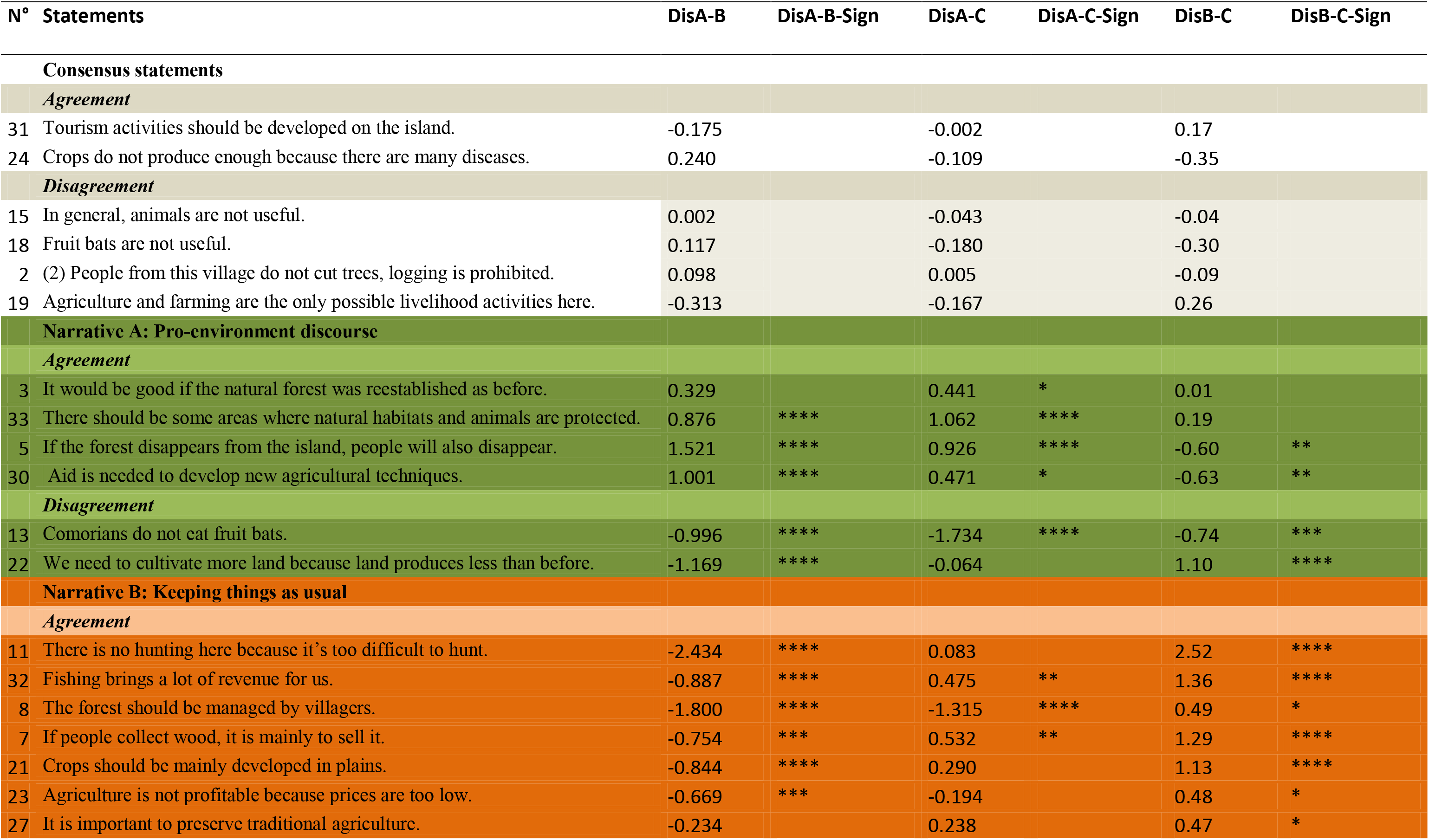
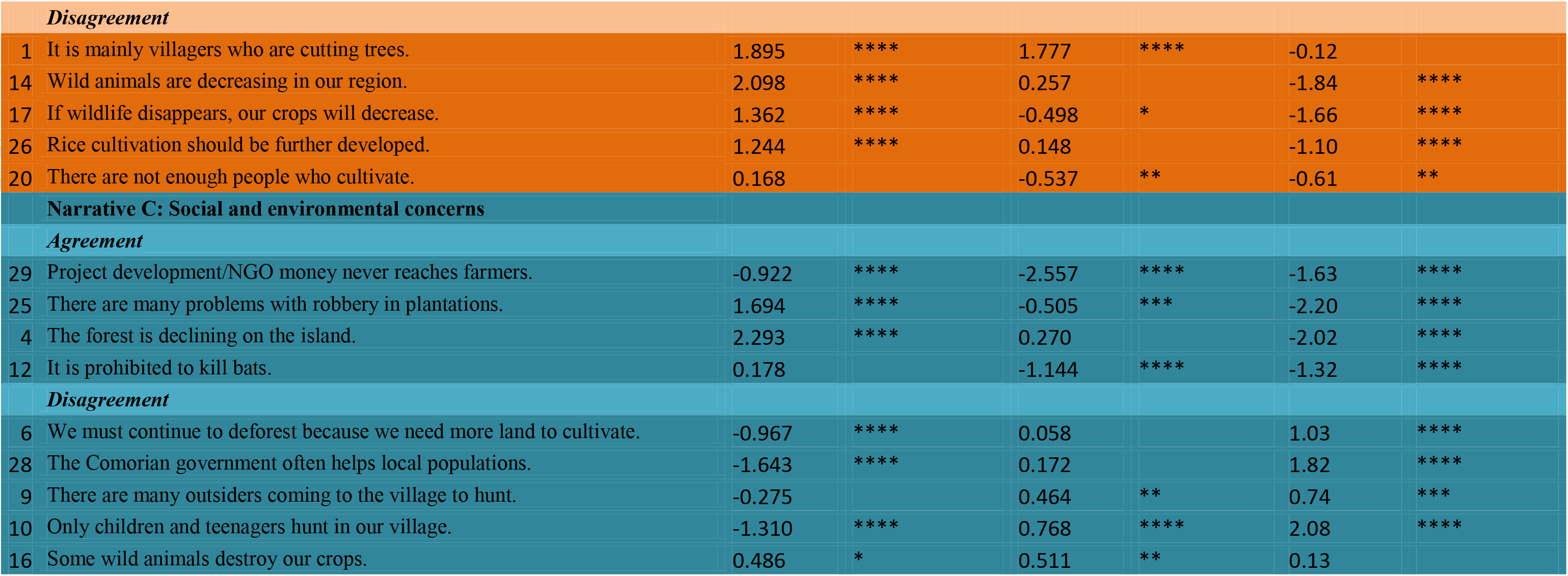
Different statements for each narrative ordered according to z-scores, beginning from the highest ranked statements in “*Agreement”* and the lowest ranked statements in “*Disagreement*”. Asterisks show statements that are distinct between narrative at P < 0.01, DisA-B=distinction between the narrative A and B; Dis A-C=distinction between the narrative A and C; Dis B-C=distinction between the narrative B and C; Sign= significantly.

#### 3.2.3 Narrative B: Keeping things as usual

Narrative B (factor 2) explained 15% of the total variance (Table s1). For this narrative, only 10 respondents loaded significantly (9 respondents from Grande Comoro and one respondent from Anjouan). These respondents were all unemployed (Unp) with a low level of education (Unp = 10 respondents). They disagreed with the statement, “It is mainly villagers who are cutting trees” [statement 1; Factor 2 score: −3], though they mostly agreed that people collect wood to sell it [statement 7; Factor 2 score: +2]. They disagreed that wild animals are decreasing in their area [statement 14]. They also disagreed that their crops may decrease if wildlife disappears [statement 17; Factor 2 score: −3]. In any case, they consider that hunting animals is too difficult and so no hunting occurs [statement 11; Factor 2 score: +4]. As one respondent who strongly disagreed (−4) commented, “It is very difficult to hunt because it requires having a gun.” Another said, “Although we would like to hunt, it is very difficult and nobody hunts here.” They believe that villagers should manage forests [statement 8; Factor 2 score: +3]. One who strongly agreed with this statement commented, “The forest belongs to the villagers and it is up to them to manage it.” Another claimed, “Forests are for villagers living nearby and who have experience in issues related to them. It is up to them to manage and to benefit from forests.” They slightly agreed that agriculture is not profitable because of low prices [statement 23; Factor 2 score: +1] and generally agreed that crops should be mainly developed in plains [statement 21; Factor 2 score: +2], but they disagreed that rice cultivation should be further developed [statement 26; Factor 2 score: −2]. They slightly agreed that it is important to preserve traditional agriculture [statement 27; Factor 2 score: +1]. They agreed that fishing brings them a lot of revenue [statement 32; Factor 2 score: +3, Table 4].

**Table 4:**
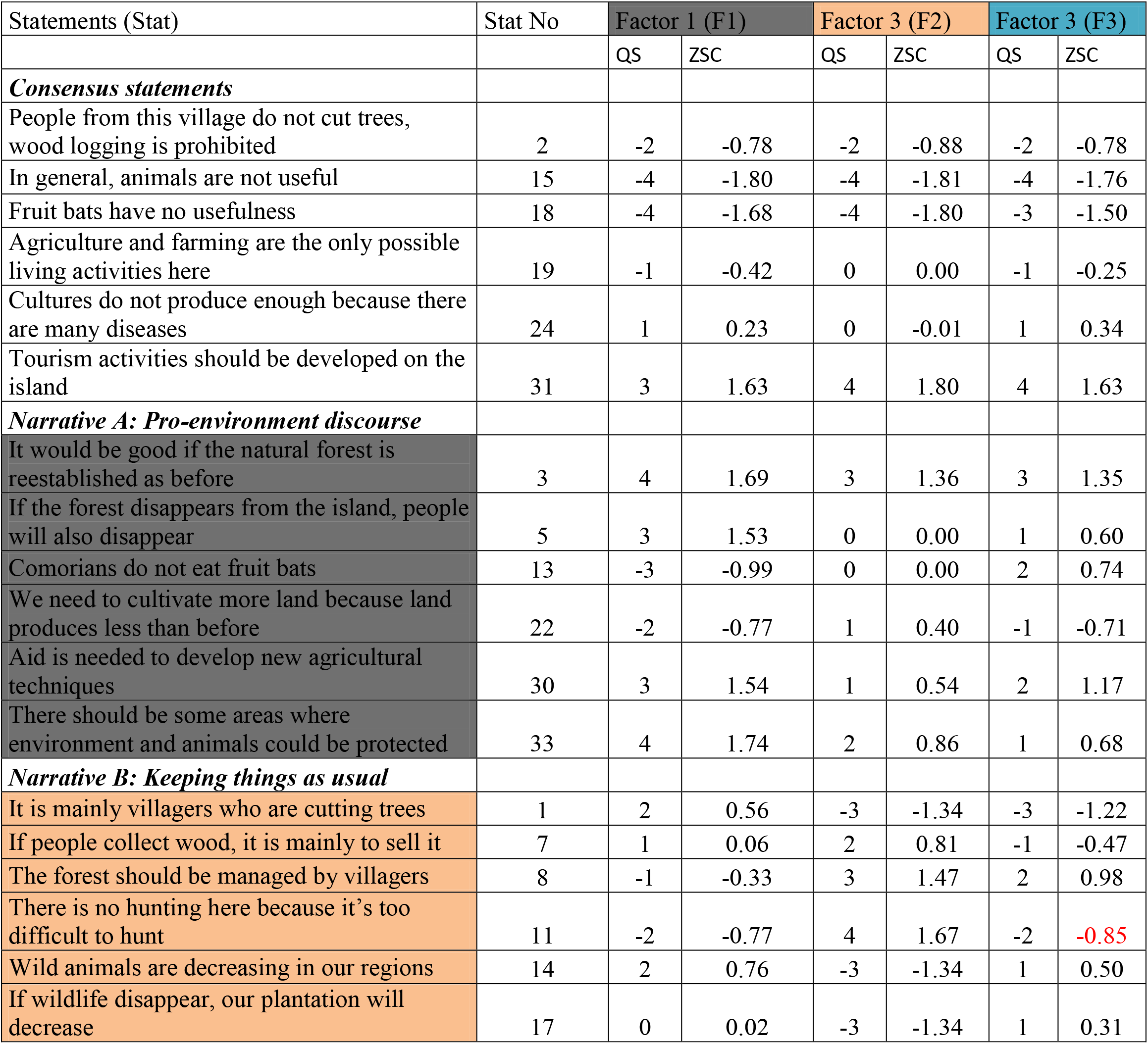
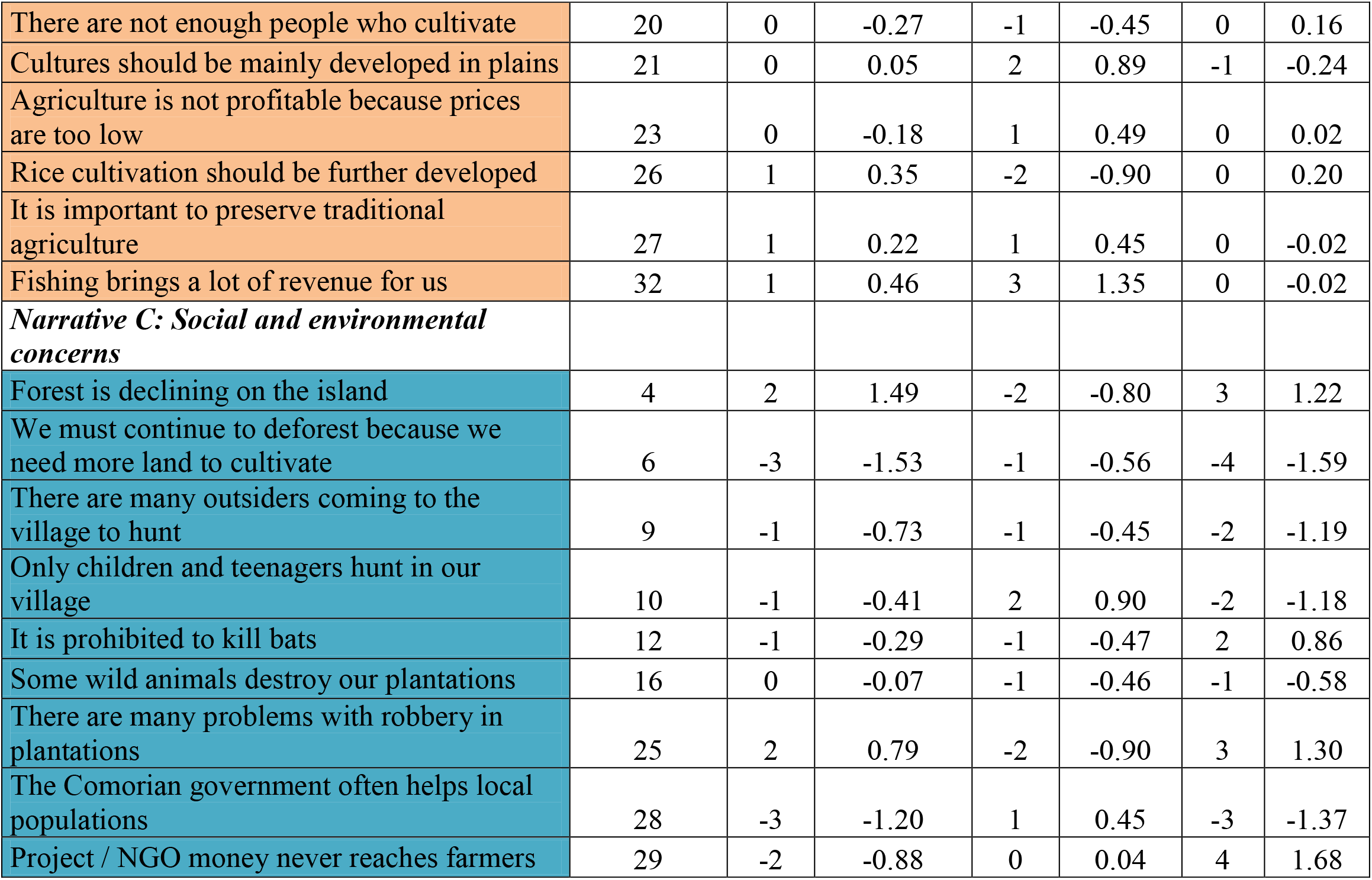
Z-scores (ZSC) and idealized Q-sort scores (QS) for the different factors or Narrative.

| Statements (Stat) | Stat No | Factor 1 (F1) |  | Factor 3 (F2) |  | Factor 3 (F3) |  |
| --- | --- | --- | --- | --- | --- | --- | --- |
|  |  | QS | ZSC | QS | ZSC | QS | ZSC |
| ***Consensus statements*** |  |  |  |  |  |  |  |
| People from this village do not cut trees, wood logging is prohibited | 2 | -2 | -0.78 | -2 | -0.88 | -2 | -0.78 |
| In general, animals are not useful | 15 | -4 | -1.80 | -4 | -1.81 | -4 | -1.76 |
| Fruit bats have no usefulness | 18 | -4 | -1.68 | -4 | -1.80 | -3 | -1.50 |
| Agriculture and farming are the only possible living activities here | 19 | -1 | -0.42 | 0 | 0.00 | -1 | -0.25 |
| Cultures do not produce enough because there are many diseases | 24 | 1 | 0.23 | 0 | -0.01 | 1 | 0.34 |
| Tourism activities should be developed on the island | 31 | 3 | 1.63 | 4 | 1.80 | 4 | 1.63 |
| ***Narrative A: Pro-environment discourse*** |  |  |  |  |  |  |  |
| It would be good if the natural forest is reestablished as before | 3 | 4 | 1.69 | 3 | 1.36 | 3 | 1.35 |
| If the forest disappears from the island, people will also disappear | 5 | 3 | 1.53 | 0 | 0.00 | 1 | 0.60 |
| Comorians do not eat fruit bats | 13 | -3 | -0.99 | 0 | 0.00 | 2 | 0.74 |
| We need to cultivate more land because land produces less than before | 22 | -2 | -0.77 | 1 | 0.40 | -1 | -0.71 |
| Aid is needed to develop new agricultural techniques | 30 | 3 | 1.54 | 1 | 0.54 | 2 | 1.17 |
| There should be some areas where environment and animals could be protected | 33 | 4 | 1.74 | 2 | 0.86 | 1 | 0.68 |
| ***Narrative B: Keeping things as usual*** |  |  |  |  |  |  |  |
| It is mainly villagers who are cutting trees | 1 | 2 | 0.56 | -3 | -1.34 | -3 | -1.22 |
| If people collect wood, it is mainly to sell it | 7 | 1 | 0.06 | 2 | 0.81 | -1 | -0.47 |
| The forest should be managed by villagers | 8 | -1 | -0.33 | 3 | 1.47 | 2 | 0.98 |
| There is no hunting here because it's too difficult to hunt | 11 | -2 | -0.77 | 4 | 1.67 | -2 | -0.85 |
| Wild animals are decreasing in our regions | 14 | 2 | 0.76 | -3 | -1.34 | 1 | 0.50 |
| If wildlife disappear, our plantation will decrease | 17 | 0 | 0.02 | -3 | -1.34 | 1 | 0.31 |
| There are not enough people who cultivate | 20 | 0 | -0.27 | -1 | -0.45 | 0 | 0.16 |
| Cultures should be mainly developed in plains | 21 | 0 | 0.05 | 2 | 0.89 | -1 | -0.24 |
| Agriculture is not profitable because prices are too low | 23 | 0 | -0.18 | 1 | 0.49 | 0 | 0.02 |
| Rice cultivation should be further developed | 26 | 1 | 0.35 | -2 | -0.90 | 0 | 0.20 |
| It is important to preserve traditional agriculture | 27 | 1 | 0.22 | 1 | 0.45 | 0 | -0.02 |
| Fishing brings a lot of revenue for us | 32 | 1 | 0.46 | 3 | 1.35 | 0 | -0.02 |
| ***Narrative C: Social and environmental concerns*** |  |  |  |  |  |  |  |
| Forest is declining on the island | 4 | 2 | 1.49 | -2 | -0.80 | 3 | 1.22 |
| We must continue to deforest because we need more land to cultivate | 6 | -3 | -1.53 | -1 | -0.56 | -4 | -1.59 |
| There are many outsiders coming to the village to hunt | 9 | -1 | -0.73 | -1 | -0.45 | -2 | -1.19 |
| Only children and teenagers hunt in our village | 10 | -1 | -0.41 | 2 | 0.90 | -2 | -1.18 |
| It is prohibited to kill bats | 12 | -1 | -0.29 | -1 | -0.47 | 2 | 0.86 |
| Some wild animals destroy our plantations | 16 | 0 | -0.07 | -1 | -0.46 | -1 | -0.58 |
| There are many problems with robbery in plantations | 25 | 2 | 0.79 | -2 | -0.90 | 3 | 1.30 |
| The Comorian government often helps local populations | 28 | -3 | -1.20 | 1 | 0.45 | -3 | -1.37 |
| Project / NGO money never reaches farmers | 29 | -2 | -0.88 | 0 | 0.04 | 4 | 1.68 |

#### 3.2.4 Narrative C: Social and environmental concerns

Narrative C (factor 3) explained 13% of the total variance. For this factor, 16 respondents loaded significantly (3 respondents from Anjouan, 6 from Moheli and 7 from Grande Comoro). These respondents were mainly unemployed with a low level of education (Unp =9), some unemployed but educated respondents (UnpE=5), while two were employed by NGOs (EmpNGO = 2), explaining the correlation between narratives A and C. They agreed with the statement that forests are declining on the island [statement 4; Factor 3 score: +3] and strongly disagreed with continuing deforestation to develop cultivated land [statement 6; Factor 3 score: −4]. As a respondent who strongly disagreed explained, “No, it is not really areas to cultivate that are lacking.”

They slightly disagreed that wild animals are destroying their crops [statement 16; Factor 3 score: −1]. They also disagreed that many outsiders come to their villages to hunt [statement 9; Factor 3 score: −2] and that only children and teenagers hunt in their village [statement 10; Factor 3 score: −2]. They agreed that it is prohibited to kill bats [statement 12; Factor 3 score: +2]. They strongly agreed that money from NGO or government projects never reaches farmers [statement 29; Factor 3 score: +4], and disagreed that the Comorian government often helps the local population [statement 28; Factor 3 score: −3]. As one respondent commented, “The Comoros government has never given assistance to local people. If it helped us, we would not be as poor as we are.” Other comments included: “The Comoros government never helps the people that is false.” “Unfortunately, NGO money is shared by agencies and does not reach the villagers.” The narrative C respondents also agreed that there are problems with robbery in cultivated areas [statement 25; Factor 3 score: +3] (Fig 3, Table 4, Table S2).”

### 3.3. Inter-class Principal Component Analysis

Considering the first three principal components, we found a high level of inter-group variation (53.40% of the total variation) between employed people (EmpNGO and Emp together) and unemployed people (Fig 4). Axis PC1 clearly differentiated between the two groups EmpNGO/Emp vs Unp, and this discrimination was significant according to the permutation test (p-value=0.01). Together, the EmpNGO and Emp groups were agreed with the following statements: “It would be good if the natural forest was reestablished as before” [statement 3], “Forests are declining on the island” [statement 4], “If the forest disappears from the island, people will also disappear” [statement 5], “Aid is needed to develop new agricultural techniques” [statement 30], “Tourism activities should be developed on the island” [statement 31], and “There should be some areas where habitats and animals are protected” [statement 33]. They were disagreed with the following statements:

**Figure 4:**
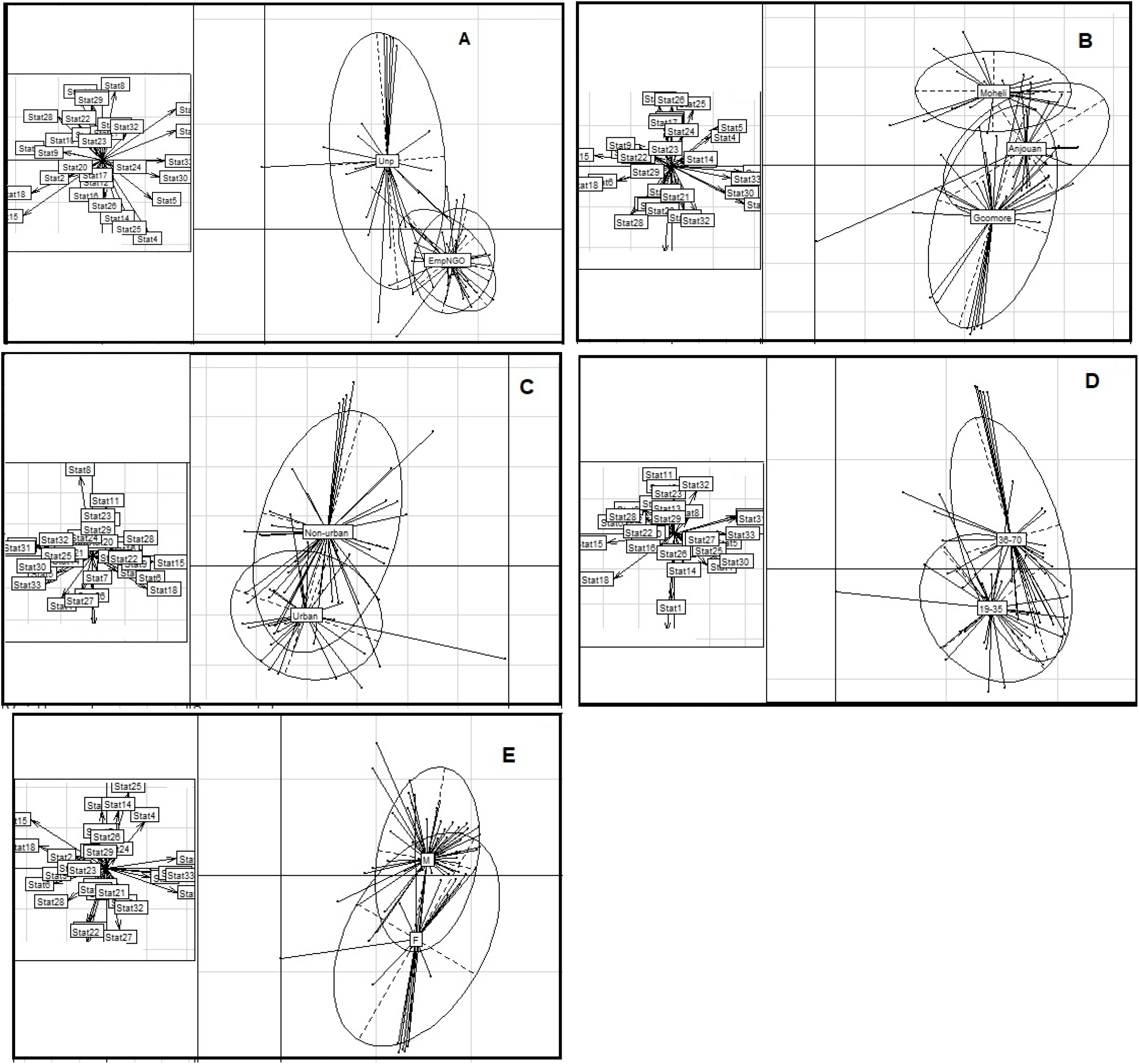
Principal component analysis (inter-class analysis). A= Discrimination between stakeholders in different social groups (EmpNGO = employed in an NGO or other; Unp = unemployed); B= Discrimination between stakeholders from the three different islands of the archipelago (Comore, Anjouan, Mohéli); C= Discrimination between stakeholders from urban and non-urban regions; D= Discrimination between Ages (from 19 to 35 years and from 36 to 70 years) of the different stakeholders; E= Discrimination between stakeholder‘s gender (M= male, F=female); The different statements are presented in left and the different groups are shown in right; the different point for each group represent the individuals; the arrows oriented to statement indicate which group the statement is more correlated on; arrows pointing up indicate that the corresponding statements are more correlated to the group at the top and arrows pointing down indicate that the corresponding statements are more correlated to the group at the bottom.

“We must continue to deforest because we need more land to cultivate” [statement 6], “Comorians do not eat fruit bats” [statement 13], “In general, animals are not useful” [statement 15], “Fruit bats have no usefulness” [statement 18], “We need to cultivate more land because the land produces less than before” [statement 22], and “The Comorian government often helps local populations” [statement 28]. The unemployed group (Unp) was positively correlated to the statements: “The forest should be managed by villagers” [statement 8], “There is no hunting here because it is too difficult to hunt” [statement 11], “We need to cultivate more land because it produces less than before” [statement 22], and “Development project/NGO money never reaches farmers” [statement 29]. It was negatively correlated to the statements: “It is mainly villagers who are cutting trees” [statement 1], “Wild animals are decreasing in our area” [statement 14], and “Rice cultivation should be further developed” [statement 26].

Considering the influence of the three islands on the first three principal components, we found a high level of inter-island variation (53.73%). Axis PC1 differentiated the three islands, and this discrimination was significant according to the permutation test (p-value =0.03). People from Grande Comore were positively correlated to the following statements: “Crops should be mainly developed in plains” [statement 21], “It is important to preserve traditional agriculture” [statement 27], and “Fishing brings a lot of revenue for us” [statement 32]. They were negatively correlated to the statements: “Agriculture and farming are the only possible livelihood activities here” [statement 19], “If wildlife disappears, our crops will decrease” [statement 17], and “Rice cultivation should be further developed” [statement 26]. People from the island of Mohéli were positively correlated to the statements: “Agriculture and farming are the only possible livelihood activities here” [statement 19], “If wildlife disappears, our crops will decrease” [statement 17], and “Rice cultivation should be further developed” [statement 26]. They were negatively correlated to the statements: “Fishing brings a lot of revenue for us” [statement 32], “It is important to preserve traditional agriculture” [statement 27], and “There are not enough people who cultivate” [statement 20]. The views of people from Anjouan were situated between those from the islands of Grande Comore and Mohéli (Fig 4).” Considering the influence of the gender, people from urban vs. non-urban regions and age classes, the discrimination test was not significant (p-value > 0.05).

## 4 DISCUSSION

### 4.1. Natural resource use by local people and its relationship to forest loss

According to the information collected in the interviews, Comorian people rely heavily on natural resources for sustenance. All (100%) of our respondents confirmed that they use the forest for cultivation or to collect wood – even those with fairly high socio-economic levels, such as administrative, financial or human resources directors. Most of the respondents have a minimum of formal knowledge about biodiversity and forests. They stated that they know what biodiversity encompasses, and they generally have a positive attitude toward wild animals. Our Q-sort sampling involved only a small number of woman‘s (23% of all respondents, see table 2.) thus our interpretation should take into account this sampling bias. The Q-sort results show that, despite the diversity of viewpoints among stakeholders, all stated the importance of forests and biodiversity, including flying fox species. However, the findings also highlight the complex links between biodiversity, natural habitats and human needs, which include the economic benefits received from agroforestry systems. Despite their understanding of the negative impacts of degraded forests on their well-being, some rural populations have no other solution for subsistence than forests and natural areas. Comorian people know that the surface area of natural habitats is decreasing in the archipelago and are aware that if the forest disappears, no human life will be possible on the islands. Most people have accurate ideas of the mechanisms involved: for instance, they detailed that complete forest loss would generate a decrease in water resources, a low yield in agriculture, a lack of charcoal and wood for building, and the disappearance of other resources, such as food, medicinal plants, etc. This indicates that Comorian people are aware of the ongoing process of degradation and its consequences, but have no alternative livelihood than to harvest in forests.

A few respondents had negative perceptions of fruit bats (raised during the interviews but not in the Q-sort surveys). These are probably due to the fact that *P. s. comorensis* feeds in cultivated areas and in fruit orchards, resulting in some damage to crops. But some respondents stated that benefits from fruit bats on their farms clearly outweigh damages. Various studies examining attitudes towards biodiversity and habitat conservation in developing countries have shown similar positive perceptions of biodiversity: for instance, in Madagascar (Ratsimbazafy et al., 2012), India (Silori, 2007; Badola et al., 2012) and Uganda (Infield and Namara, 2001). In our study, positive perceptions of biodiversity were largely driven by the perceived benefits to the respondents. For example, most positive attitudes toward *P. livingstonii* were due to the fact that the species attracts many tourists as it is one of the largest bats as well as one of the most threatened animals in the world, but also because the species plays a crucial role in forest regeneration and in crop cultivation. The positive attitudes toward *P. s. comorensis* were related to its role as a seed disperser, but also to the fact that the species represents an important source of food for many rural populations.

Our results identified three main discourses, or narratives, one of which (Pro-environment discourse) supports long-term biodiversity conservation through the creation of protected areas. This narrative recognizes the consequences of forest loss and supports the development of ecological agricultural methods that allow forests to be maintained and developed. The second narrative (Keeping things as usual) is more in favor of immediate benefits from the forest and the protection of local activities and revenues, despite the awareness of the importance of forests and the effects of natural habitat loss on local livelihoods. The third narrative (Social and environmental concerns) is in favor of immediate benefits from forests, but equally sees the necessity of preserving natural habitats. These respondents understand the importance of preserving forests and the negative impact of forest and biodiversity loss, but are forced by poverty to harvest natural resources. Positive attitudes toward long-term biodiversity conservation (Pro-environment discourse) are held mainly by employed people, including NGO staff, professors, agricultural engineers and other public officials. This could be linked to the fact that their employment leads them to be less dependent on forests and natural resources. Many previous studies have shown a significant relationship between employment, formal education and perceptions of biodiversity and forest conservation (King and Peralvo, 2010; Cairns et al., 2013).

Our results indicated that respondents with a low level of formal education, who are often unemployed, are associated with narrative “Keeping things as usual”. Being dependent on forest resources, their main concern is to protect their livelihoods rather than biodiversity leading them to stress that only local people should manage forests and natural resources. This highlights that the lack of other means of securing the necessities of life is the main factor leading rural people to harvest natural resources. While they are aware of the broad importance of forests, for these people, protecting them is essential mainly for their subsistence or health rather than for intrinsic or ecological reasons.

According to our analysis, the Narrative B appears to represent an attitude associated to Grande Comores respondents (10 respondents loaded significantly among which 9 respondents from Grande Comoro and one respondent from Anjouan). The results must be interpreted with caution as may not be broadly applicable across the three islands of the Union of Comoros.

Among unemployed respondents, five were educated (level of university), among which three have just finished the university studies and two of them are finished education since few years but do not have formal jobs. These unemployed but educated respondents mainly belonged to the narrative (Social and environmental concerns, see table 2) indicating that they are aware of the necessity of preserving natural habitats but are also in favor of immediate benefits from forests because of the level of poverty in these islands. Despite their high education level and their awareness regarding the importance of forests, biodiversity and natural habitat, these people are poor and struggle to meet their day-to-day needs and are in favor of any actions that may generate immediate benefits for their survival. This highlights that although education is crucial for understanding and awareness regarding the importance of forest and biodiversity conservation, reducing poverty and increasing livelihoods of local people of these islands is the key strategy to allow habitat and biodiversity conservation actions to be effective.” These rural people claimed that aid money never reaches farmers. In the Comoros, development project budgets are often managed by people with a high level of education, and local people believe that this money is always absorbed by these agencies. As aid from development projects and NGOs is often limited, and thus insufficient to reach all rural people, this leads those who do not benefit to have a negative perception of NGOs.

Our results found that rural people from Grande Comore and Anjouan intensively collect wood to sell it, resulting in a high harvesting rate of the forests of these islands compared to Mohéli forests (Granek et al., 2002; Sewall et al., 2011, Ibouroi et al., 2018a). In contrast, respondents from Mohéli are in favor of forest and biodiversity conservation, including the development of ecological rather than traditional agriculture (the latter is preferred by respondents from Grande Comore). Respondents from Mohéli feel that wood collection should be prohibited in their region. On this island, due to the presence of the National Park of Mohéli, various nature conservation projects, and the high level of tourism linked with local biodiversity (e.g. sea turtles, Livingstone‘s flying fox etc.), biodiversity represents the main source of income for the population (Granek and Brown, 2005).

Our study‘s findings highlight the diversity of viewpoints among Comoros stakeholders depending on several social factors, including formal education level, employment, and geographic location. These results join a number of other studies that have shown diverse local perceptions of biodiversity and how to manage natural resources (Watkins and Cruz, 2007; Tapia et al., 2009a; Gall and Rodwell, 2016; Kamal and Grodzinska-Jurczak, 2014). Understanding the nuances in attitudes and the different weights attributed by stakeholders to each element of the dilemma may help to find unexpected areas of agreement and to advance new solutions.

### 4.2. Conservation recommendations

Previous studies in the Comoro Islands have proposed different strategies for limiting intensive forest exploitation including law enforcement, deployment of the national army in forest, educational initiatives such as increasing awareness and understanding of conservation issues (Mikuš, 2009; Poonian et al.2008a; Trewhella et al., 2005). However, all these strategies do not appear as appropriate solutions for effectively reducing habitat destruction since Comorians’ exploitation of natural resources is a question of survival. As many stakeholders commented during our interviews, “We use natural resources for our survival. We will continue to exploit forests even if it costs our life.” In the other hand, our results indicate that Comorians today do not lack awareness concerning the importance of natural habitats and the impact of habitat disturbance and loss on their livelihood. Rather it appears that the main constraint is poverty, forcing them to heavily exploit forests. In addition, employing force as a conservation measure is dangerous for villagers, forest managers and conservationists. Some Mohéli respondents affirmed that marine turtle poachers are often armed. In an assessment of Comorians’ perception of the Mohéli Marine Park, a Marine Protected Area (MPA), Poonian et al. (2008b) revealed that the most important factors affecting habitat management in the protected areas of Comoros are the lack of sustainable alternative livelihoods, inequitable distribution of benefits and continuing environmental threats. Poonian et al. (2008b) suggested that, to ensure habitat conservation and the continuity of this protected area, MPA managers should adopt programs that carefully consider sustainable sources of finance for stakeholders and lower-cost alternatives that reduce poverty. Hauzer et al. (2008) highlighted that Comorians, especially from Mohéli, were aware of the importance of the protected area, but felt that their survival was of priority importance. Hauzer et al. (2008) suggested that the best conservation strategy would be a measure that would “(1) ensure sustainability through effective financial planning and appropriate management techniques; (2) mobilize local communities to create a truly co-managed MPA; (3) ensure tangible benefits to local communities through realistic alternative livelihood options”. In a study of the links between resource dependency and attitude of commercial fishers to coral reef conservation in the red sea, Marshall et al. (2010) found a direct relationship between conservation attitudes and aspects of resource dependency. Especially, fishers with higher income were more likely to have a positive conservation attitude. Sewall et al. (2011) suggested that local Comorians living near forests should be compensated if agricultural land use within a reserve were restricted. One of the most important management strategy in protected area is involving local people and habitat users in the management (Mtwana et al 2014). Freed and Granek (2014) suggested that priority for management actions should be to include local community members and stakeholders in the decision-making and implementation process for protecting fragile reef ecosystems in the Comoros. These authors suggested that local communities would serve as the primary management actor for an effective conservation strategy (see also Freed et al 2016). Sewall et al. (2011) also suggested that any plans for a reserve should be adopted through a formal process that includes local community engagement, as without this conservation strategies will not be effective.

Our results highlight that employment influence local perceptions and suggest the lack of livelihoods for rural people as the main factor leading local people to overharvest natural resources. Although these rural people cultivate more, most of them apply traditional methods especially by using slash-and-burn making lands unproductive few years letter and increasing the need of more lands.

The first key recommendation we propose for the preservation natural habitats is developing and maintaining sustainable production of crops for local human benefit. New methods and materials to develop ecological agriculture must be made available to local communities. Rocliffe et al. (2014) highlighted that underdevelopment of legal structures supportive of local communities was one strong constraints for formal local protected areas in many developing countries of the Western Indian Ocean islands including the Comoros archipelago. Projects such as these could allow local populations to improve yield with the same surface area, thus reducing the conversion of forest into farmland.

The lack of market to sell cultivated products is the second factor leading to overexploitation of natural habitats as highlighted by many respondents. As a second key recommendation we propose to develop projects of local markets that that could allow the creation of new jobs for local people.

According to our results (semi-structured interviews), all interviewed respondents are using forests and natural resource for subsistence. Fisher and Christopher (2007) highlighted that, about 72% of Comorians depend directly on forest resources. This strong dependence on natural resources is due to the fact that many development sectors such as tourism are not yet developed in the Union of Comoros (Granek and Brown, 2005). The third key recommendation we propose is to develop eco-touristic project, including the construction of bungalows in strategic villages as well as tourist sites for observing emblematic species such as the endemic flying foxes, lemurs, scops owl, etc. Villagers and local communities could manage these infrastructures.

As the lack of governmental assistance is claimed to be the main cause leading to the overharvesting of forests, the fourth key recommendation we propose here is that government aids and support should made available for rural people that could reassure them of the government good intentions to contribute to local development.

The fifth key strategy allowing ensuring the preservation of Comoros forests and natural habitats in mid-term and long-term purposes will be to set up awareness campaign for replantation and reforestation. As Comorian people do not lack awareness regarding the necessity to preserve forests and natural habitats but overexploit forest for their everyday needs, forest managers must ensure the improvement of living conditions of local population before any replantation project otherwise community will exploit the forests before replanted trees grow. Replantation can play an important role in sustaining native biodiversity and makes an important contribution to the conservation of native biodiversity (Rocliffe et al. 2014). Their re-establishment involves the replacement of native natural plants but also makes large trees available for mid and long-term purposes which are crucial for human wellbeing (Brockerhoff et al. 2008).

## 5. CONCLUSION

In conclusion, habitat loss and the vulnerability of biodiversity in the Comoros are the results of the unsustainable overexploitation of natural resources. Yet to maintain the ecological balance necessary for daily human needs and for future generations (clean water, productive agricultural land, ecosystem services from biodiversity and forests), it is vital to conserve the natural habitats on these islands.

As the exploitation of natural resources by local people is a question of survival, a program allowing to reduce poverty for instance by developing tourism and maintaining sustainable production of crops, livestock and setting up awareness campaign for tree planting and reforestation projects are necessary for the Union of Comoros.

On the three islands of the Union of Comoros, a project to create national marine and terrestrial protected areas – including national parks funded by the Global Environmental Finance (GEF) and put in place since 2016 by the United Nation Development Program (UNDP) – has been agreed by the Comorian government. The project is now managed by an independent institution (The National Network of Protected Areas or *Réseau National d’Aires Protégées RNAP*). Based on our interviews with local people, most rural communities agree with the creation of protected areas if they can gain direct benefits from the project and are involved in the conservation actions. Constructive engagement with local residents (such as providing employment as local guides or park rangers, for example) would contribute to supporting long-term conservation success.

## Acknowledgments

We would like to acknowledge the Direction of Environment and Forest of Comoros for giving permission to carry out our field works (Permit N° 002/KM/15/DNEF). Field work was funded by the Rufford Foundation through a Research Support Grants (Grant N° 21803-1 to SAOD and N° 26731-2 to MTI).

